# GlucoPush: A Do-It-Yourself Add-On for Online Tracking of Personal Glucometer Use

**DOI:** 10.1101/296293

**Authors:** Luis R Soenksen, Zulkayda Mamat

**Affiliations:** L. R. Soenksen is with the Mechanical Engineering department at the Massachusetts Institute of Technology, Cambridge, MA 02139 USA.; Z. Mamat is with Biological Engineering department at the Massachusetts Institute of Technology, Cambridge, MA 02139 USA.

**Keywords:** Connectivity, device, diabetes, do-it-yourself, glucometer, internet-of-things, point-of-care, wireless

## Abstract

Current public health guidelines on diabetes management recommend frequent self-monitoring and physician tracking of blood glucose to reduce complications caused by this condition. While Internet-of-Things glucometers exist to aid in this goal, most commercial glucometers seen in wide-spread use worldwide do not provide these online tracking capabilities or allow for convenient data sharing with health care providers. This situation is caused, among several factors, due to premium pricing of IoT glucometers and resistance from older users to dispose of their previous glucometers. We propose an add-on strategy to enable IoT integration in glucometers to increase the use of wireless connectivity features in this sector. Here we describe and test a simple do-it-yourself system fabricated with low-cost commercially available wireless connectivity components that aim to demonstrate this IoT augmentation strategy to track glucose readings from previously unconnected standard glucometers.

## I. INTRODUCTION

According to the 2016 Global Report on Diabetes by the World Health Organization, the prevalence of diabetes in the world has nearly doubled since 1980 with an estimated 422 million adults currently living with this chronic disease [1]. With this number on the rise, the advent of systems enabling better self-monitoring of blood glucose (SMBG) has become increasingly relevant towards the goal of reducing complications and improving quality of life. Current clinical guidelines recommend frequent SMBG for most diabetic patients to guide insulin control therapy, physical activity and food intake [2]. Moreover, reliable tracking of SMBG records is now considered an essential step towards empowering patients and health care providers who benefit from reduced hospitalizations caused by diabetes-related complications [3]. Online data tracking has already been integrated into a broad range of commercially available glucometers, a feature that is now easily implementable to a range of systems due to the recent introduction of low-cost wireless connectivity and Internet-Of-Things (IoT) modules. Despite their availability, IoT glucometers are still sold as premium products, with price-points starting at around $40 as compared to low-cost standard options starting at $8 [4]. This 5-fold increase in price causes purchasing hesitation in many customers who could otherwise benefit from more personalized diabetes management enabled by online tracking of glucose measurements. This specific pricing strategy has complex origins, but follows from regulatory barriers for new medical products, protection of profitable product pipelines owned by key manufacturers, and various costumer preferences. Moreover, for many diabetic patients who have already invested in a specific glucometer model with months’ worth of strip supplies, the idea of transitioning to a different IoT-enabled device may be perceived as wasteful. Therefore, it appears that developing an add-on strategy to enable IoT integration of previously unconnected glucometers could facilitate the widespread use this connectivity feature. Here we present a simple do-it-yourself system fabricated with low-cost commercially available wireless connectivity components that aims to demonstrate a simple and low-cost IoT augmentation strategy to track glucose readings from standard but previously unconnected glucometers. With this, we aim to inform product designers, entrepreneurs, patients and physicians on the feasibility of simple approaches towards the development of add-on commercial solutions for glucometers and other point-of-care diagnostics based on this concept to address current connectivity needs.

## II. METHOD OF APPROACH AND DESIGN

### A. Overall System

The GlucoPush glucometer tracking add-on system (Fig. 1) consists of several components – (1) A film pressure sensor which is used to detect the insertion of a blood glucose test strip into the commercial glucometer, (2) a low-cost WiFi module that pushes notifications to a user’s personal cell phone via open local networks, (3) a secure mobile web application accessible through the push notification to incentivize the user to report their glucose level reading and fill a short questionnaire to share with their doctor.

**Fig. 1.**
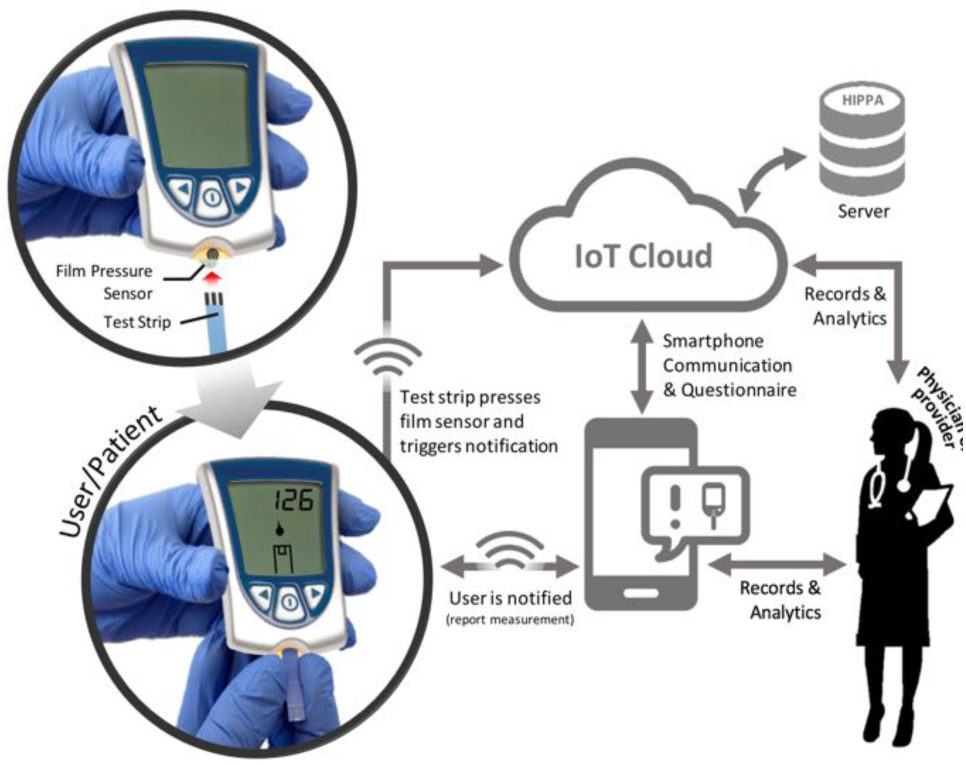
Schematic of the overall functionality of the GlucoPush add-on system. Insertion of the test strip in the glucometer compresses the installed film pressure sensor which then triggers a notification for measurement storage in secure cloud via a smartphone.

### B. Film Pressure Trigger

To detect the insertion of the blood glucose test strip, a force sensitive flex-film resistor (FSFFR) with a 0.16” (4 mm) diameter active sensing area (Interlink Electronics, CA) is used. The active area of the FSFFR sensor is placed at the test strip entry port to detect strip insertion. The resistance of the FSFFR sensor varies depending on the level of pressure applied to the sensing area. A threshold of pressure indicating full strip insertion can be defined or calibrated depending on the geometric tolerances of the glucometer in use. The two 0.1-inch pitch terminals of the FSFFR sensor extend from the bottom of the sensing area to a soldering connection to allow compatibility with Arduino boards and common electronic components. This FSFFR sensor was installed at the strip port of a Precision Xtra^TM^ Glucometer (Abbott Laboratories Inc., Lake Bluff, IL) as shown in Fig. 2A. The average film pressure sensor resistance without strip insertion was around 1MΩ, whereas the average resistance under film compression due to full test strip insertion was approximately 2.5kΩ. A voltage divider using Vref = 3.0V is defined using this sensor and an additional 1MΩ reference resistor, which is then interrogated using an analog-to-digital converter (ADC) port in the system microcontroller. A voltage threshold of Vref/2 maintained for >10 sec was selected to indicate a strip insertion event.

**Fig. 2.**
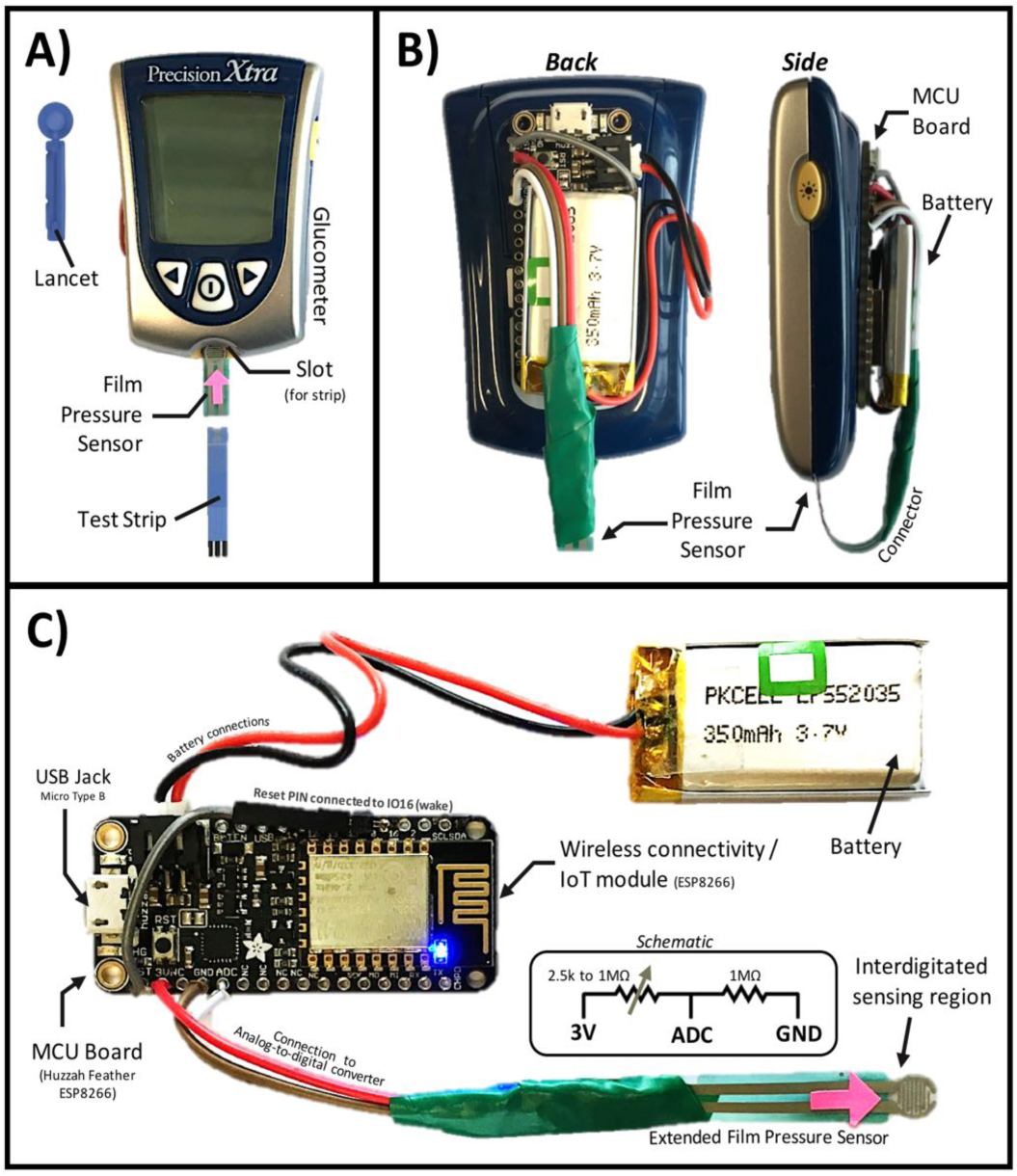
Fabrication and installation detail of GlucoPush add-on. A) Front view of a commercially available glucometer system with installed pressure sensitive film in test strip slot. B) Back and side views of installed GlucoPush add-on in a commercial glucometer. C) Detail of add-on components, showing the microcontroller unit (MCU) and IoT module, as well as rechargeable battery and film pressure sensor.

### C. Microcontroller and Firmware

The GlucoPush system is controlled by an Adafruit HUZZAH Feather (Adafruit, NY), which is a WiFi and Bluetooth-enabled board with an ESP8266 microcontroller (80-160 MHz with 3.3V logic/power) and built-in USB for programming and charging. This light-weight board (approx. 6 grams), shown in Fig. 2B and 2C, was selected due to its robust auto-reset functions, built-in connector for a rechargeable 3.7V 350mAh Lithium polymer battery (Adafruit, NY), as well as its popularity for open-source IoT applications. The ESP9266 microcontroller in the Huzzah board was programmed to provide the desired IoT glucometer capabilities using code provided in the <u>supplemental material</u> and the Arduino IDE. The programmed firmware includes a WiFi connection routine that automatically detects open networks at the user’s location, and a notification routine that sends a secure link to a pre-defined patient’s smartphone without the use of a dedicated cellular data plan or a short message service (SMS). The encrypted notification is pushed to the user’s personal smartphone via an application called Pushover (Superblock, IL) as shown in Fig. 3. The compilation-ready “.INO” file for Arduino IDE of the system is included in the <u>supplemental material</u>, and just requires user’s modification with a personal encryption key and Pushover account.

**Fig. 3.**
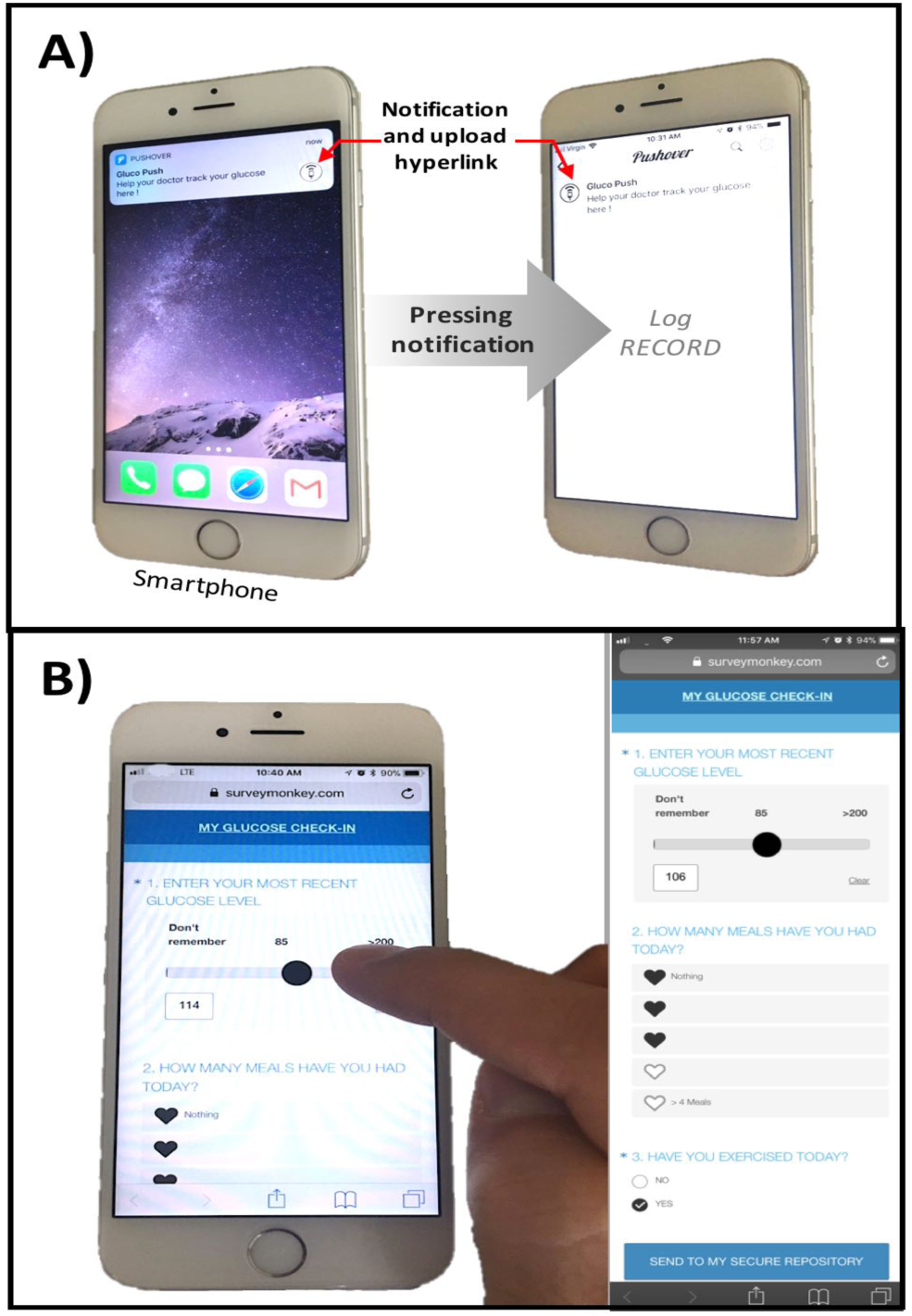
GlucoPush notification and measurement input system. A) Shows notification and log. B) Snapshot of smartphone input online survey for glucose levels recording. The collected data can be shared to provider via a secure HIPPA compliant connection.

### D. Mobile Application

Following the standard measurement workflow of the used glucometer, the user is automatically prompted with a push notification on their personal smartphone as seen in Fig. 3A. This notification is achieved using a Pushover framework, which is a free to download app and needs to be previously installed in the user’s smartphone. The notification contains an active hyperlink to a short online questionnaire made using SurveyMonkey (SurveyMonkey, CA) as shown in Fig. 3.

This online recording system can be modified to use other web-accessible links or systems and is not limited to the presented questionnaire. To provide secure operation, the GlucoPush system is programmed with a key chosen by the user/provider, this same key is then used by the Pushover application to push notifications to the user’s smartphone. The mobile questionnaire platform allows the glucose reading data to be permanently stored and shared by the user at their own discretion since they are the only ones with access to the same key.

## III. USABILITY TESTING

A general usability test was conducted with three independent users (age 50±17) to evaluate the ease of operating the add-on in a commercially available glucometer. The users were given minimal instruction for installation of hardware and software, as well as general operation of the system. User’s capacity to use the device once installed and programmed was assessed, as well their ability to assemble and program the device on their own.

## IV. RESULTS AND DISCUSSION

The GlucoPush system was fabricated for under $10 only using off-the-shelf and open-source components. All evaluated users were able to adequately use the GlucoPush add-on device when it had been previously installed in the glucometer of their choosing and was also appropriately linked to their own smartphone device. Only one user with previous electrical engineering and programming experience was able to assemble the device, as well as install the required firmware and software without help. This suggests that the GlucoPush add-on (at least in its current form) requires experience in circuit assembly and programming. Furthermore, basic familiarity with smartphone and Pushover mobile app seems to also be needed to effectively access the questionnaire sent with the pushed notification.

## V. CONCLUSION

The proposed add-on prototype cannot yet be considered an off-the-shelf solution and is only intended to allow for user experimentation in this do-it-yourself approach for IoT integration of glucometers. To our knowledge, this is the first time such add-on strategy has been implemented to connect standard glucometers in a way that allows for a patient-centered control over the stored and/or shared glucose monitoring information. We hope our design triggers further innovation and allows for a natural and much-needed cooperation between developers, patients with diabetes and physicians. We recognize that a study with a representative number of diabetic users is still needed to fully evaluate usability and clinical benefit of our strategy. A particularly important group to serve and analyze with such new SMBG systems includes older populations, who become particularly attached to specific glucometer brands or user interfaces. Since new devices may hold unknown modes of operation and failure, these patients are likely to hesitate when given the opportunity to switch or participate in an unknown IoT-enabled system. Such barriers for adoption are relevant, even in the context of informed and price insensitive diabetic patients that recognize the importance of constantly sharing actionable information with their physicians. Despite the simplicity of our design, replicating our prototype requires prior electrical engineering and programming experience that is not accessible to most target users. Therefore, we hope this design allows other developers to refine our IoT integration methodology into an easy-to-use and secure off-the-shelf product to share glucose information with health care providers with the objective of better personal management of diabetes.

## ACKNOWLEDGMENT

This project was funded and supported by the Harvard–MIT Program of Health Sciences and Technology, Department of Mechanical Engineering, and Department of Biological Engineering at Massachusetts Institute of Technology (MIT).

